# Unique amino acid substitution in RBD region of SARS-CoV-2 Omicron XAY.2

**DOI:** 10.1101/2023.01.09.523246

**Authors:** Lekha Salsekar, Shefali Rahangdale, Ekant Tamboli, Krishna Khairnar

## Abstract

We attempted to explain the rare mutation at the receptor binding domain of the spike protein in the XAY.2 variant of SARS-CoV-2 from the perspective of hydrophobic interactions. We propose that decreasing hydrophobicity at position 446 and 486 of the RBD region of the spike protein might affect the infectivity of SARS-CoV-2. We also estimated the probable mutations at the 446 and 486 position the virus may acquire, leading to a decreased hydrophobicity.

## BACKGROUND

XAY.2 is an emerging omicron variant detected mainly in Denmark 77.0%, South Africa 15.0%, and Israel 8.0% [1]. It was also detected recently in Thailand. This variant is of concern due to two unique mutations, G446D and F486P, in the receptor binding domain (RBD) region of the SARS-CoV-2 spike protein. G446D and F486P are shown to decrease and increase infectivity, respectively; this could also affect the efficacy of spike protein-based SARS-CoV-2 vaccines [2, 3]. The Spike protein plays a significant role in the attachment of the SARS-CoV-2 virus to the host cell receptors. The spike protein’s RBD residues, in particular, form the RBD-Human Angiotensin Converting Enzyme (hACE) complex, which facilitates membrane fusion and virus entry into human cells. Therefore, any modification in the RBD is crucial in terms of the fitter virus variant [4,5].

### SIGNIFICANCE OF SPIKE PROTEIN MUTATION IN RBD AT POSITIONS 446 & 486

Positions 446 and 486 of the spike protein, along with others, are pivotal positions in terms of binding of RBD to hACE2 [2,6]. The omicron sublineages have been observed to be prone to acquire mutations at positions 446 and 486; the mutational analysis revealed a unique mutation, G446D, in XAY.2 when compared to 16 other omicron sublineages which have G446S mutation (Figure). Another rare and unique mutation of F486P was observed in XAY.2 and also XBB.1.5, which is due to two nucleotide polymorphism in the same codon when compared to the wild-type Wuhan strain [1]. BA.2.75.2, XBB, XBB.1, XBB.2, and CH.1.1 have F486S mutation; and BA.4, BA.5, BA.4.6, BQ.1, BQ.1.1, and BF.7 have F486V mutation (Figure).

**Figure legend:**
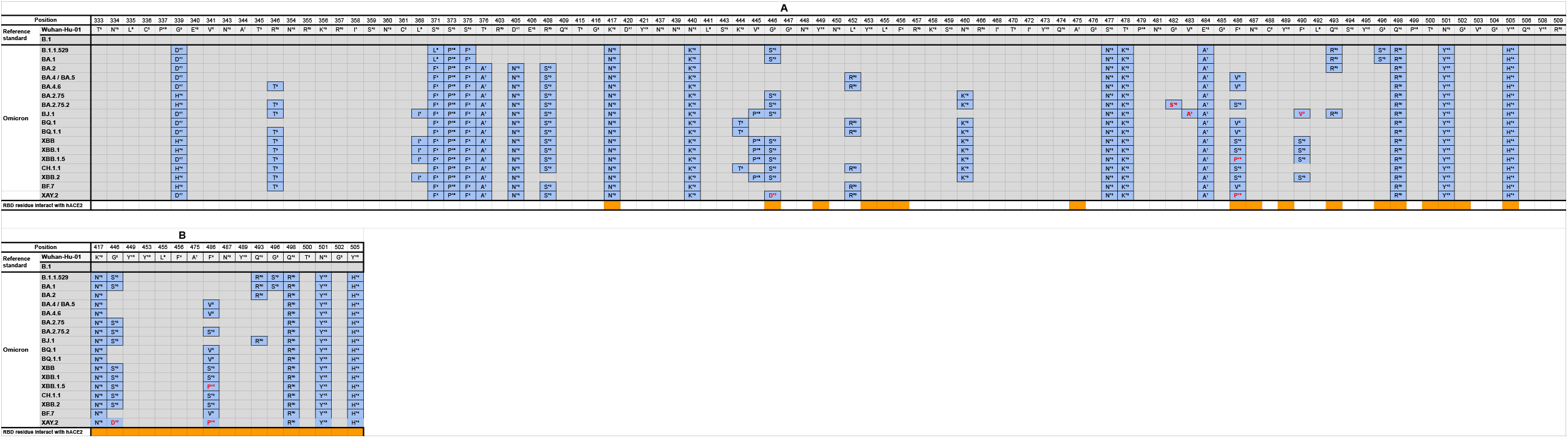
(A) A comparison of 17 globally most prevalent Omicron sublineages for their respective positions in the RBD region. Mutations in the RBD of Omicron sublineages. Superscripts denote relative hydrophobicity on a scale of 1 to 20 (from higher to lower hydrophobicity) of amino acids according to Kyte and Doolittle scale [7]. Significant novel mutations to the omicron sublineages are highlighted in red, and the position for RBD residue interacting with hACE2 is indicated beneath the sequences in orange. (B) A comparison of Omicron sublineages for only RBD residues interacting with hACE2. RBD residues interacting with hACE2 have been identified for 17 positions. Significant novel mutations to the omicron sublineages are highlighted in red, and the position for RBD residue interacting with hACE2 is indicated beneath the sequences in orange.

### THE HYDROPHOBICITY OF AMINO ACIDS AFFECTS THE BINDING

The mutational history of omicron sublineages, particularly XAY.2, shows that the selection pressure pushes it to acquire the amino acid residues that are relatively less hydrophobic than the wild-type Wuhan strain. In the XAY.2, the amino acids at spike 446 and 486 positions are aspartate and proline, which are less hydrophobic than the wild-type amino acids glycine and phenylalanine, respectively [7]. A summary of amino acid hydrophobicity is denoted on a scale of 1 to 20 (higher to lower) as per the study of Kyte and Doolittle [7], which has been represented in the Table.

**Table:** Amino acid substitutions at position 486 and 446 observed in the RBD region of Spike for 17 Omicron Sublineages.

| Amino acids In RBD region of spike protein in omicron sublineage | 446 Position of Spike Protein<br>BA.1, BA.2.75, BA.2.75.2, BJ.1, XBB, XBB.1, XBB.1.5, CH.1.1 and XBB.2 have G446S mutation; and XAY.2 have G446D | 486 Position of Spike Protein<br>BA.2.75.2, XBB, XBB.1, XBB.2, and CH.1.1 have F486S mutation; and BA.4, BA.5, BA.4.6, BQ.1, BQ.1.1, and BF.7 have F486V | Possible Substitutions at in emerging variants |  |
| --- | --- | --- | --- | --- |
|  |  |  | 446 | 486 |
| I<sup>1</sup> ATT, ATC, ATA |  |  |  |  |
| V<sup>2</sup> GTT, GTC, GTA, GTG |  | V<sup>2</sup> GTT, GTC, GTA, GTG |  |  |
| L<sup>3</sup> CTT, CTC, CTA, CTG, TTA, TTG |  |  |  |  |
| F<sup>4</sup> TTT, TTC (Wild type for position 486) |  |  |  |  |
| A<sup>7</sup> GCT, GCC, GCA, GCG |  |  |  |  |
| G<sup>8</sup> GTC, GAC |  |  |  |  |
| T<sup>9</sup> ACT, ACC, ACA, ACG |  |  |  |  |
| S<sup>10</sup> TCT, TCC, TCA, TCG, AGT, AGC | S<sup>10</sup> TCT, TCC, TCA, TCG, AGT, AGC | S<sup>10</sup> TCT, TCC, TCA, TCG, AGT, AGC |  |  |
| Y<sup>12</sup> TAT, TAC |  |  |  |  |
| P<sup>13</sup> CCT, CCC, CCA, CCG |  | P<sup>13</sup> CCT, CCC, CCA, CCG |  |  |
| H<sup>14</sup> CAT, CAC |  |  |  | H<sup>14</sup> CAT, CAC |
| D<sup>17</sup> GAT, GAC | D<sup>17</sup> GAT, GAC |  |  | D<sup>17</sup> GAT, GAC |
|  |  |  | 446 | 486 |
| N<sup>18</sup> AAT, AAC |  |  | N<sup>18</sup> AAT, AAC | N<sup>18</sup> AAT, AAC |
| K<sup>19</sup> AAA, AAG |  |  | K<sup>19</sup> AAA, AAG | K<sup>19</sup> AAA, AAG |
| R<sup>20</sup> CGT, CGC, CGA, CGG, AGA, AGG |  |  | R<sup>20</sup> CGT, CGC, CGA, CGG, AGA, AGG | R<sup>20</sup> CGT, CGC, CGA, CGG, AGA, AGG |

For a surface-interacting protein like the spike protein, the phenomenon of hydrophobicity plays a significant role in affecting its interaction with the receptor [8]. When the highly hydrophobic and neutral phenylalanine amino acid is substituted by a less hydrophobic and neutral proline amino acid at the 486 position of spike protein, this increases RBD-hACE2 binding affinity. However, at position 446, the highly hydrophobic and neutral glycine amino acid is substituted with a less hydrophobic and negatively charged aspartate amino acid; this decreases RBD-hACE2 binding affinity. So far, by this discussion, one can presume that having a higher degree of hydrophobicity of amino acid is unfavourable for the protein to interact with another protein in an aqueous environment. However, this is not entirely the case; hydrophobic residues also help stabilise the protein in an aqueous environment by forming a stable core. Studies revealed that these hydrophobic residues also stabilise protein-ligand binding [9]. However, some hydrophobic pockets are also on the surface, which repels the water molecules away from the binding site and makes the binding site residues more available to form hydrogen bonds with the ligand preventing the binding site from getting saturated with water molecules [10]. It is important to note that substituting an amino acid that has decreased hydrophobicity with a neutral charge increases RBD-hACE2 binding affinity. Substitution with an amino acid that has decreased hydrophobicity with negative charge decreases RBD-hACE2 binding affinity.

### SELECTION PRESSURE DRIVEN AMINO ACID SUBSTITUTION THAT ARE LESS HYDROPHOBIC

We can observe a trend of decreasing hydrophobicity of amino acids for the mutations at position 486 of spike protein in omicron sublineages. Amino acids having lower hydrophobicity are getting selected. We propose that if the 486 position continues to incorporate amino acid substitutions which are less hydrophobic, then the probability of substitutions including His, Asp, Asn, Lys, and Arg is more. Out of these five probable amino acid substitutions, except Lys, all other amino acid substitutions are likely to occur by substituting two nucleotide bases in the spike gene at position 486 compared to the wild type. Further, we also propose that if the 446 position continues to incorporate amino acid substitutions which are less hydrophobic, then the probability of substitutions including Asn, Lys, and Arg is more.

## References

1. Lineage List. https://cov-lineages.org/lineage_list.html (accessed Jan 09, 2023).

2. Verma J, Subbarao N. Insilico study on the effect of SARS-CoV-2 RBD hotspot mutants’ interaction with ACE2 to understand the binding affinity and stability. Virology. 2021 Sep;561:107–116. doi: 10.1016/j.virol.2021.06.009. Epub 2021 Jun 28. PMID: 34217923; PMCID: PMC8237243.

3. Wang, Q., Iketani, S., Li, Z., Liu, L., Guo, Y., Huang, Y., Bowen, A. D., Liu, M., Wang, M., Yu, J., Valdez, R., Lauring, A. S., Sheng, Z., Wang, H. H., Gordon, A., Liu, L., & Ho, D. D. (2022). Alarming antibody evasion properties of rising SARS-CoV-2 BQ and XBB subvariants. Cell. https://doi.org/10.1016/j.cell.2022.12.018

4. Jackson, C. B., Farzan, M., Chen, B., & Choe, H. (2021). Mechanisms of SARS-CoV-2 entry into cells. Nature Reviews Molecular Cell Biology, 23(1), 3–20. https://doi.org/10.1038/s41580-021-00418-x

5. Ding, C., He, J., Zhang, X., Jiang, C., Sun, Y., Zhang, Y., Chen, Q., He, H., Li, W., Xie, J., Liu, Z., & Gao, Y. (2021). Crucial Mutations of Spike Protein on SARS-CoV-2 Evolved to Variant Strains Escaping Neutralization of Convalescent Plasmas and RBD-Specific Monoclonal Antibodies. Frontiers in Immunology, 12. https://doi.org/10.3389/fimmu.2021.693775

6. Shang, J., Ye, G., Shi, K., Wan, Y., Luo, C., Aihara, H., Geng, Q., Auerbach, A., & Li, F. (2020). Structural basis of receptor recognition by SARS-CoV-2. Nature, 581(7807), 221–224. https://doi.org/10.1038/s41586-020-2179-y

7. Kyte, J., & Doolittle, R. F. (1982). A simple method for displaying the hydropathic character of a protein. Journal of Molecular Biology, 157(1), 105–132. https://doi.org/10.1016/0022-2836(82)90515-0

8. Young, L., Jernigan, R., & Covell, D. (1994). A role for surface hydrophobicity in protein-protein recognition. Protein Science, 3(5), 717–729. https://doi.org/10.1002/pro.5560030501

9. Sarkar, A., & Kellogg, G. (2010). Hydrophobicity - Shake Flasks, Protein Folding and Drug Discovery. Current Topics in Medicinal Chemistry, 10(1), 67–83. https://doi.org/10.2174/156802610790232233

10. Li, J., Ma, X., Guo, S., Hou, C., Shi, L., Zhang, H., Zheng, B., Liao, C., Yang, L., Ye, L., & He, X. (2020). A Hydrophobic-Interaction-Based Mechanism Triggers Docking between the SARS-CoV-2 Spike and Angiotensin-Converting Enzyme 2. Global Challenges, 4(12), 2000067. https://doi.org/10.1002/gch2.202000067

